# Plant resistance to drought relies on early stomatal closure

**DOI:** 10.1101/099531

**Authors:** N Martin-StPaul, S Delzon, H Cochard

## Abstract

Plant resistance to drought has long been thought to be associated with the ability to maintain transpiration and photosynthesis longer during drought, through the opening of stomata. This premise is at the root of most current framework used to assess drought impacts on land plants in vegetation models. We examined this premise by coupling a meta-analysis of functional traits of stomatal response to drought (i.e. the water potential causing stomatal closure, *Ψ*_close_) and embolism resistance (the water potential at the onset of embolism formation, *Ψ*_12_), with simulations from a soil-plant hydraulic model. We found that *Ψ*_close_ and *Ψ*_12_ were equal (isometric) only for a restricted number of species, but as Ψ12 decreases, the departure from isometry increases, with stomatal closure occurring far before embolism occurs. For the most drought resistant species (*Ψ*_12_<-4.2 MPa), *Ψ*_close_ was remarkably independent of embolism resistance and remained above −4.5 MPa, suggesting the existence of a restrictive boundary by which stomata closure must occur. This pattern was supported by model simulations. Indeed, coordinated decrease in both *Ψ*_close_ and *Ψ*_12_ leads to unsuspected accelerated death under drought for embolism resistant species, in contradiction with observations from drought mortality experiments. Overall our results highlight that most species have similarity in stomatal behavior, and are highly conservative in terms of their water use during drought. The modelling framework presented here provides a baseline to simulate the temporal dynamic leading to mortality under drought by accounting for multiple, measurable traits.

## INTRODUCTION

Recent drought episodes have been identified as the triggers for widespread plant mortality events around the world [1–3]. They have had huge consequences for the productivity of the land [4] and have undoubtedly affected a panel of ecosystem services [5]. Identifying the mechanisms and traits underlying drought resistance is essential to understanding and predicting the impact of widespread droughts over many land areas. Experimental studies have provided empirical evidence that the failure of the water transport system is tightly linked to tree dehydration and mortality in drought conditions. This was confirmed by a recent study reporting that global patterns of mortality was predictable from hydraulic safety margins[6]. Two key types of traits shaping the trade-off between drought resistance and the maximization of carbon dioxide assimilation have been identified: hydraulic traits ensuring the integrity of the hydraulic system under drought [7], and stomatal traits controlling gas exchanges at the leaf surface [8]. However, efforts to model tree mortality in response to drought is still hindered by a lack of understanding of how these traits interact to define physiological dysfunctions under drought stress [9]. In this study, we analyzed the overall connections between these two types of traits for the full range of drought resistance, with a soil-plant hydraulic model.

The stomata have two key functions: controlling transpiration, which supplies nutrients and regulates leaf temperature, and controlling the entry of CO_2_ into the leaf. Stomatal closure in response to water deficit is the primary limitation to photosynthesis [10], and constitutes a key cost in terms of plant growth and leaf temperature under drought conditions. However, stomatal closure also limits excessive decreases in water potential (quantified as a negative pressure, *ψ*) in the plant, thereby ensuring that the water demand from the leaves does not exceed the supply capacity of the hydraulic system, which would lead to embolism of the vascular system and complete dehydration of the plant. These key, but opposing roles of stomata in regulating CO_2_ influx and H_2_O loss pose a dilemma that has occupied scientists for centuries^1^ and has led to the view that plant stomata probably operate at the edge of the supply capacity of the plant’s hydraulic system, to balance different cost such as productivity leaf temperature regulation during drought [11,12].

Conversely, maintenance of the supply capacity of the hydraulic system depends on the ability of a species to sustain high negative pressure to limit embolism. Embolism resistance is usually quantified by the value of *ψ* causing 50% embolism (*Ψ*_50_), and the rate of embolism spread per unit drop in water potential (*slope*). From these two traits the *Ψ* at the onset embolism formation can be computed (*Ψ*_12_, see equations 1 to 3), which give a more conservative functional limit to the hydraulic system. Embolism resistance varies considerably between species and with the dryness of species habitat [7,13,14]. A recent study has suggested that hydraulic systems highly resistant to embolism have evolved in response to the selective pressure associated with increasing drought levels during a paleoclimatic crisis [15]. Some contemporary plants have extremely drought-resistant vascular systems, with *Ψ*_50_ values reaching −19 MPa [16].

These findings have led to the suggestion that an efficient match between the capacity of the hydraulic system to sustain water deficit (*i.e.* embolism resistance) and the regulation of demand by the stomata is a prerequisite for the maximization of gas exchanges without dehydration [11,17–19]. This notion naturally leads to the hypothesis that stomatal behavior and embolism resistance have followed a similar evolutionary trajectory under drought constraints, and that plants have increased their intrinsic embolism resistance to allow stomata to close later during drought, thereby maximizing plant productivity [8,12,20]. The coordination of stomatal and hydraulic traits and their role in shaping drought resistance has yet to be addressed on a global scale. This would help to clarify the interplay between mechanisms and plant traits in defining the physiological dysfunctions occurring under drought stress, which remains one of the principal challenge faced in the modeling of tree mortality in response to drought.

In this study we gathered stomatal regulation traits and embolism resistance traits for different species. We compiled *Ψ*_12_, *Ψ*_50_ and *slope* values derived from stem vulnerability curves published by us during the past 20 years, representing 151 species from different biomes (Tables S1). Recent direct observations of embolism formation by mean of X-ray tomography [21–24] confirmed the reliability of these values. For those species, we performed a seach for data of water potential causing stomatal closure (*Ψ*_close_). We used concurrent measurements of gas exchange and leaf water potential, from which the *Ψ* value at 90% stomatal closure was calculated following [8,20]. Stomatal opening increases with guard cell turgor pressure [25–27], thus, we also used leaf water potential at turgor loss (*Ψ_tl_*) as a surrogate for *Ψ*_close_. We then explored the range of variation and the coordination between *Ψ*_close_ and *Ψ*_50_. Finally, we used a soil-plant water transport model to elucidate how different associations between *Ψ*_close_ and *Ψ*_50_ determine the time until hydraulic failure during drought (see Methods). We validated model predictions using empirical data for time to shoots death collected in drought mortality experiments[28–31] (see Methods and Appendix 1).

## RESULTS AND DISCUSSION

Embolism resistance (taken as the *Ψ*_50_) ranged between −1.3 and −19 MPa (Figure 1 A). The large variations of *Ψ*_50_ were partly related to changes of the *slope* which was non linearly related to the *Ψ*_50_ (Figure 1 A, insert). Both the *Ψ*_50_ and *slope* determined the value of the water potential causing the onset of embolism (*Ψ*_12_, equation 3). This more conservative indicator of embolism resistance ranged between −0.7 and −14 MPa. The two indicators of water potential causing stomatal closure (*Ψ*_close_) were significantly related between each other with a slope close to one as previously reported [20] (p.value<0.01 Figure 1b insert). *Ψ*_close_ was thus taken as the average value when the two traits were found. By contrast to embolism resistance, *Ψ*_close_ varied from −1. to −4.3MPa, spanning a three times lower range of variation than embolism resistance (Figure 1 A & B), in agreement with recent meta-analysis [20].

**Figure 1:**
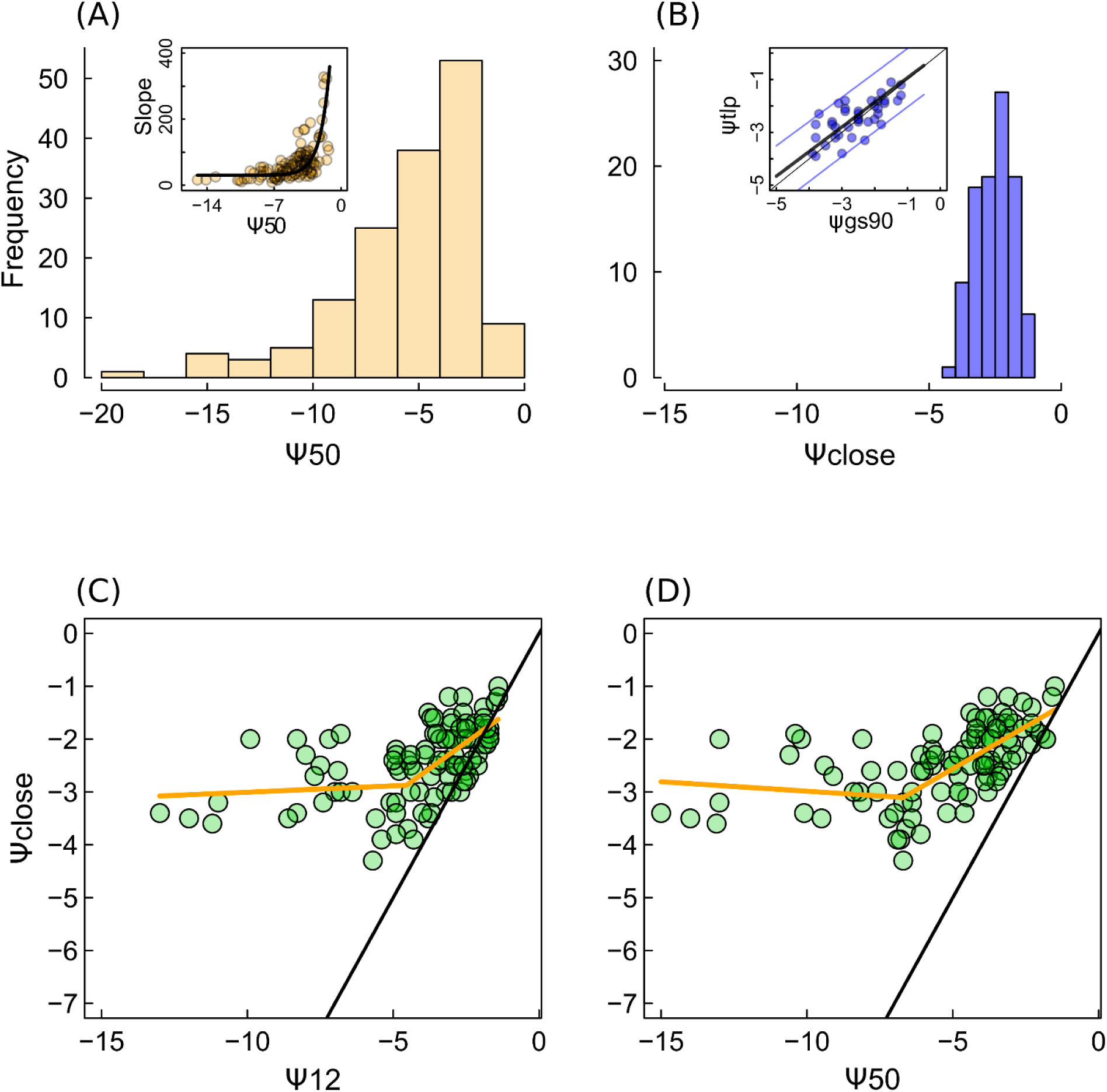
Range of variation of embolism resistance (*Ψ*_50_) and stomatal response to drought (*Ψ*_close_), and their covariation, showing that these two traits do not covary isometrically. (A) Distribution of embolism resistance (*Ψ*_50_) among plants. The insert shows the relationship between the *slope* of the vulnerability curve (%/MPa, equation 2) and *Ψ*_50_, that allows to compute *Ψ*_12_ (MPa). The best fit was obtained with *slope*= 16+e^(*Ψ*_50_)^×1092, the fitted parameters were significatly different from 0 (P.value<0.01). (B) Distribution of the water potential causing stomatal closure (*Ψ*_close_). The relationship between the two different traits for *Ψ*_close_ (the water potential causing 90% stomatal closure, *Ψ*_gs90_) and the water potential causing leaf turgor loss (*Ψ*_tlp_) is shown in the inset with fitted line (*Ψ*_tlp_=0.97×*Ψ*_gs90_, p.value<0.01, R^2^=0.4) and the 95% confidence interval. (C) The relationship between *Ψ*_close_ and *Ψ*_12_, and (D) relationship between *Ψ*_close_ and *Ψ*_50_. Points are individual species and the orange line is the best fit with a segmented regression, showing a significant break point for *Ψ*_50_ of −5.9 MPa or *Ψ*_12_ of −4.2MPa corresponding to an average *Ψ*_close_ of ca. −3 MPa (± 1.5MPa, 95%CI).

The relationship between *Ψ*_12_ and *Ψ*_close_ did not follow the isometric line (i.e. the 1:1 line, Figure 1c), but had a slope lower than unity: 0.4 for species with low embolism resistance ( *Ψ*_12_>-4.2) and null for drought resistant species (For *Ψ*_12_<-4.2, p.value=0.4). This contradicts the expected match between *Ψ*_12_ and *Ψ*_close_ and indicates that the departure between *Ψ*_12_ and *Ψ*_close_ increase with increasing embolism resistance. These results were confirmed when we used *Ψ*_50_ as an indicator of embolism resistance (Figure 1 D, Table S2). The relationship between *Ψ*_close_ and embolism resistance (*Ψ*_50_, *Ψ*_12_) presented a marked interruption (i.e. for *Ψ*_12_ of −4.2 or *Ψ*_50_ ca.-6 MPa) corresponding to *Ψ*_close_ =-3Mpa on average (Figure 1 C, Figure 1 D and Table S2). Overall, these results indicate that stomatal closure always occurs at much higher water potential value than the one triggering embolism and that the difference between *Ψ*_close_ and *Ψ*_50_ increases continuously with increasing embolism resistance. Contrasting the hypothesized coordination between stomatal closure and embolism resistance, our finding supports a similarity in stomatal behavior particularly among embolism resistant species, and indicates that most species are highly conservative in terms of their water use during drought.

The similarity of stomatal behavior between species suggests that keeping stomata open is not beneficial in terms of fitness, particularly for survival under drought conditions. To get more insights into the relationship between stomatal function and plant resilience to drought, we developped a soil plant-hydraulic model (see *Methods*) computing the survival times under drought conditions (i.e. the time to hydraulic failure) for the range of hypothetical species covering the full spectrum of embolism resistance. We used three different postulates to assess stomatal behavior (Figure 2 A, see *Methods* and Appendix 2). Firstly (hypothesis 1), we assumed that stomata do not close to regulate transpiration (*E*) during drought (i.e. stomata are maintained open whatever the soil and plant water potential). Secondly, we assumed (hypothesis 2) that stomata regulate water losses to maintain plant water potential (*Ψ* plant) above the water potential resulting in the onset of embolism (i.e. *Ψ*_12_), according to the premise that the integrity of the hydraulic system is functionally linked to stomatal closure in response to drought. This was done by reducing *E* to a residual term (*E*_min_) when *Ψ*_plant_ approached *Ψ*_12_ (Appendix 4). Finally (hypothesis 3), we assumed that *Ψ*_close_ varied with *Ψ*_50_ until −3 MPa, independently of the hydraulic properties of the vascular system (orange line in Figure 2c), in accordance with the empirical average trend observed in our dataset (Figure 1 C). We simulated the *E* decline due to the progressive loss of turgor as water deficit increases, by inverting the pressure volume curve equations[32] (Appendix 3). At turgor loss point, *E* was reduced to *E*_min_, as for hypothesis 2. We report on Figure 2b the relationships between survial time (time to hydraulic failure) and *Ψ*_50_ modeled for each of the three hypothesis. As a mean of validation, we compared these relationships with results from drought mortality experiments in which survival time increased with *Ψ*_50_ (Figure 2 C).

**Figure 2:**
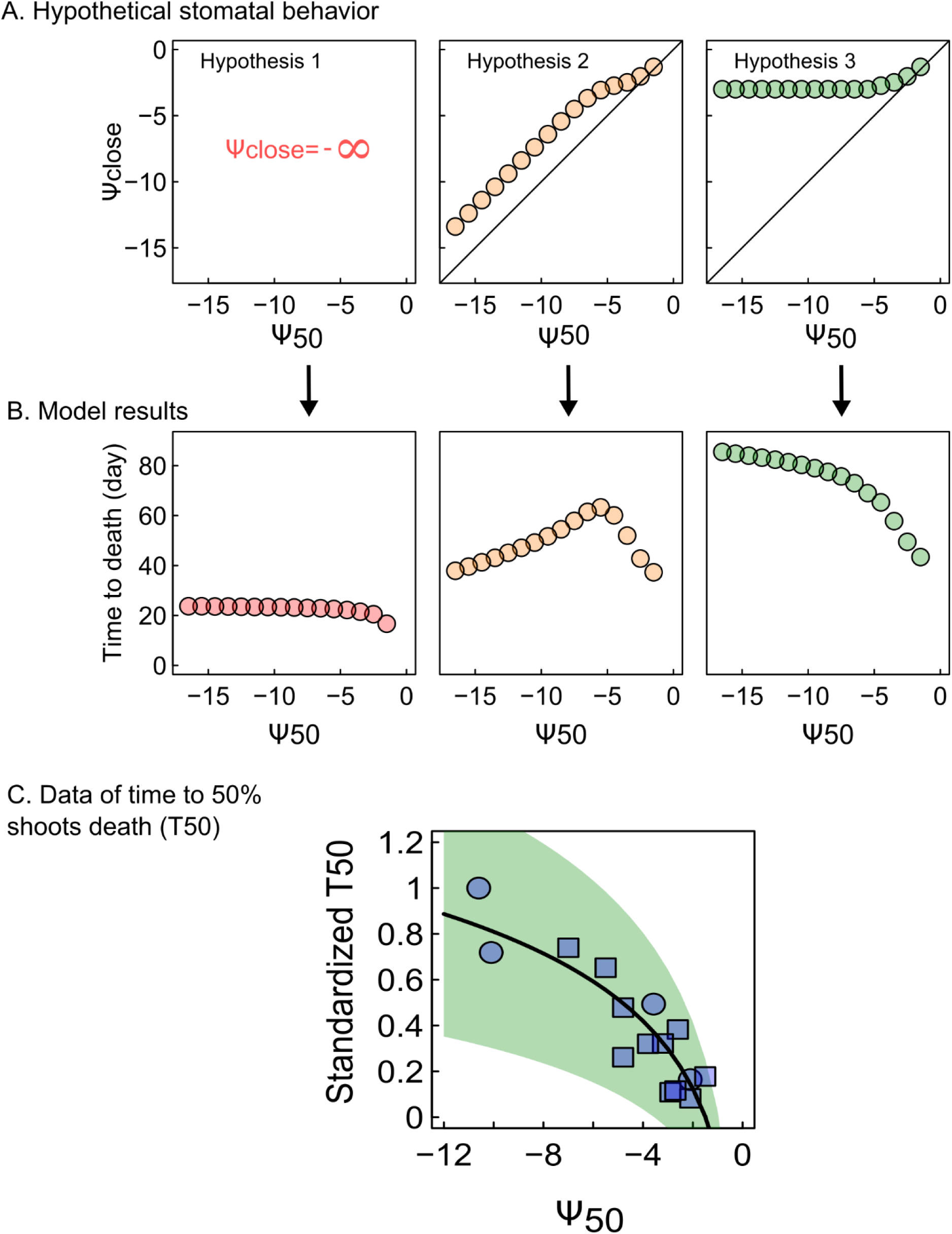
Model simulations of the survival time for the range of embolism resistance, under three different hypothesis of stomata behaviour. (A) Representation of the parameters combinations for *Ψ*_close_ and *Ψ*_50_ used to represent the three hypothesis tested in the model: (Hypothesis 1) Stomata never close *(i.e.* plants maintain maximal rates of transpiration at all soil and plant water potentials whatever their *Ψ*_50_). (hypothesis 2) Stomata regulate water losses to maintain plant water potential above the water potential resulting in the onset of embolism (*i.e* stomata close when water potential reaches *Ψ*_12_). (hypothesis 3) Stomata closure and embolism resistance were not coordinated untill −3MPa as indicated by our empirical results (Figure 1c,d). (B) Simulated relationship between survival time (time until hydraulic failure) and *Ψ*_50_ for each hypothesis tested. (C) Normalized time to 50% shoot death (T_50_) as a function of *Ψ*_50_ for 15 species. The data were gathered from four different studies and normalized to account for differences in soil and climate conditions across experiments (see *Methods).* The logarithmic relationship fitted on absolute values (0.42×log(|*Ψ*_50_|)-0.16, slope p.val<0.001) is shown (line) with 95% CI (green area).

Hypotheses 1 and 2 were not consistent with the empirical observation that survival time increased with embolism resistance (Figure 2 C). By contrast, simulations imposing early stomatal closure (hypothesis 3) resulted in an increase in survival with embolism resistance (Figure 2 B).

Under hypothesis 1, plants would die very rapidly from hydraulic failure, with only a slight increase in survival when *Ψ*_50_ decreased from −1 to −6 MPa (Figure 2 C). For *Ψ*_50_ values below −6 MPa, increasing embolism resistance was not associated with a further increase in survival time, suggesting that increasing embolism resistance *per se* had only a marginal impact on survival under drought conditions. If water loss regulation was imposed to maintain *Ψ*_plant_ just above the level triggering embolism (hypothesis 2), survival time was much higher (Figure 2 B), and increased markedly with embolism resistance until *Ψ*_50_ reached −6 MPa. However, beyond this value, survival decreased substantially (Figure 2B), contrary to the trend observed in analyses of experimental induced mortality (Figure 2B). This increase in survival for at high *Ψ*_50_ (*Ψ*_50_>-6MPa), is in fact due to the maintenance of *Ψ*_12_ (and thus of stomatal closure) at a constant of ca. −3 MPa because of the *slope* of the VC decrease with decreasing *Ψ*_50_ (Appendix A2-4, Figure A2-6). If the *slope* of the VC was maintained at a constant average value of 45%/MPa for all the *Ψ*_50_ tested, the model would simulate a consistant decrease of survival with embolism resistance under this assumption (Appendix 2-2, Figure A2-2).

The survival peak simulated at a *Ψ*_50_ of −6 MPa (Figure 2 B) under this hypothesis, implies that *Ψ*_close_ is about −3 MPa, corresponding to the mean limit for *Ψ*_close_ in our database. Consequently, assuming that *Ψ*_close_ does not covary with embolism resistance beyond *Ψ*_close_ = −3 MPa (hypothesis 3), a positive relationship between survival and embolism resistance was predicted over the entire range of *Ψ*_50_ (Figure 2 B), consistent with the empirical trend observed in drought mortality experiments (Figure 2 A). These simulations provided support for the view that embolism resistance cannot increase survival unless the difference between embolism resistance and *Ψ*_close_ also increases.

An analysis of the modeled dynamics of soil and plant dehydration for two species with contrasting level of embolism resistance identified the physical mechanisms making early stomata closure a prerequisite for the avoidance of drought-induced mortality, even for embolism-resistant species (Figure 3). The relationship between soil water potential (*Ψ*_soil_) (and, hence, plant water potential) and soil water content (Θ) becomes nonlinear at relatively high value of *Ψ*_soil_ (Figure 3a and 3b). Thus, the longer transpiration is maintained the sharpest are the rates of soil and plant water potential drops, leading to rapid death through hydraulic failure. The nonlinearity of the *Ψ*_soil_ (Θ) relationship results from long-established physical laws [33,34] describing the changes in *Ψ*_soil_ and soil conductivity with soil water content. These laws are globally conserved among soil types (Appendix A4, Figure A4-1), providing support for the global scope of our findings.

**Figure 3:**
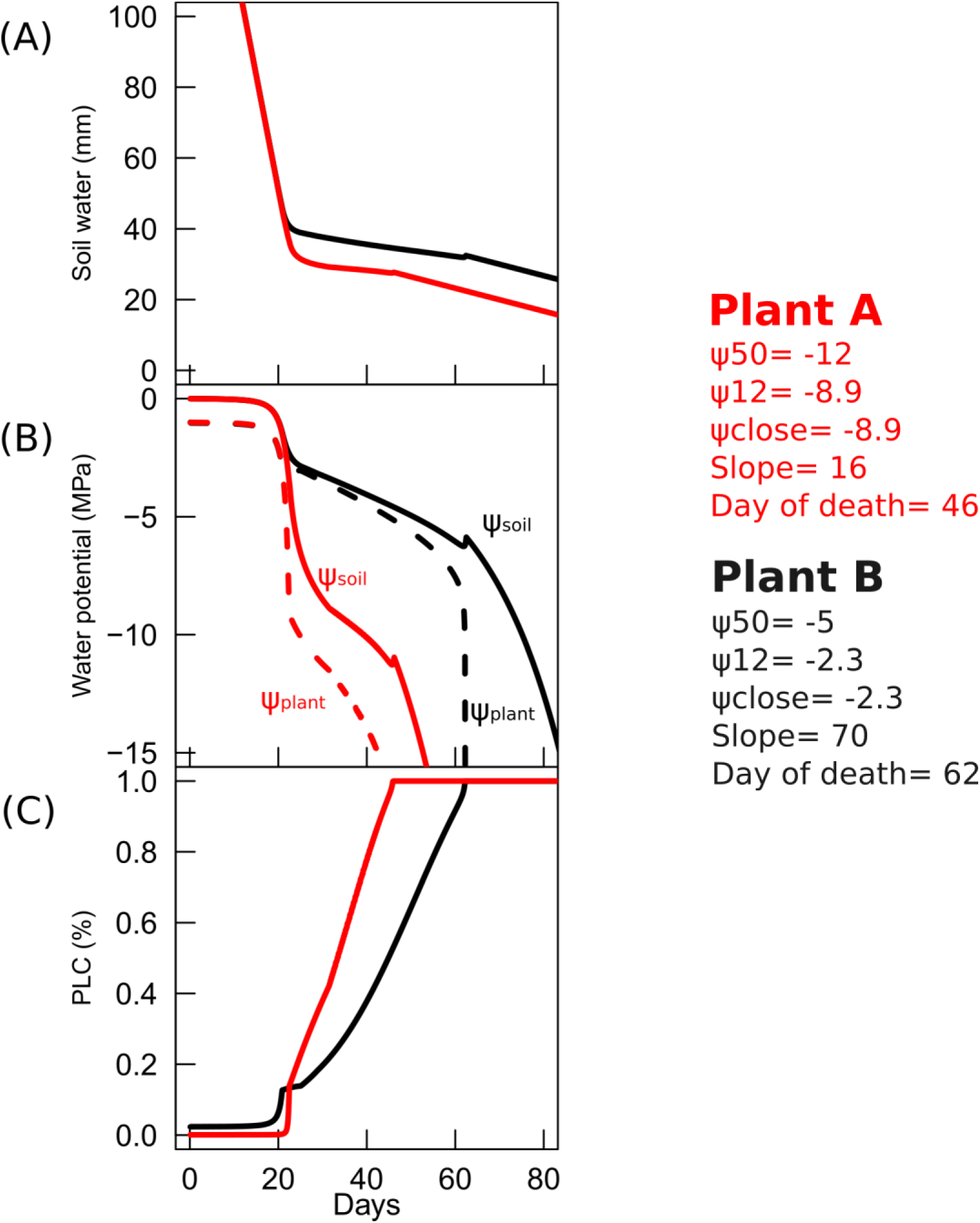
Simulated temporal dynamics of soil and plant dehydration assuming that stomata close when water potential reaches *Ψ*_12_ (hypothesis 2 in the text) for two contrasted species. (A) soil water content, (B) soil and plant water potential and (C) the percent loss of conductivity caused by embolism. Simulations were performed for two hypothetic plants (plant A and plant B) which traits are given in MPa on the plot. The time to death by hydraulic failure (i.e. 100% embolism) is also indicated. Simulations show that high levels of embolism resistance, and thus stomatal closure at lower water potential accelerated death, because of faster water potential drops. More detailed simulation results are given Appendix 2.

The vascular system of terrestrial plants has evolved toward very high levels of embolism resistance (*Ψ*_50_ values down to −19 MPa) allowing the colonization of dry environments^21^. Stomatal closure was thought to have evolved along similar lines, to keep carbon assimilation levels for longer periods, even at low xylem water potential. Different recent studies have moreover reported tight co-variations between stomatal closure to drought and embolism resistance, but for relatively low drought resistant species [8,12,20]. Our results highlight that the range of variation of *Ψ*_close_ appears much reduced when seen in the light of the full range of embolism resistance. Such uncoupling between stomata closure and the failure of the vascular system may be the result of selection pressures that have favored survival under extreme water scarcity over growth under mild drought.

These findings provide a complementary view to the widely accepted framework for drought response strategies based on the water to carbon trade-off that plants have to face(e.g[35][36]). According to this framework, plant drought response strategies fall in between two extreme categories called isohydric and anisohydric [8,35,37,38]. Isohydric plants close their stomata rapidly in response to drought, thereby maintaining a high water potential to limit embolism, but at the risk of death due to carbon starvation. Conversely, anisohydric plants keep their stomata open at low water potential, maintaining carbon assimilation levels, but at the cost of damage to the water transport system through embolism. This framework has been the focus of many scientific studies on drought-induced mortality in recent decades and has underpinned our understanding and modeling of drought induced plant mortality [35,36,39]. The fact that plants among the most drought resistant close their stomata at much higher potential than embolism can occur, indicates that resisting drought may not involve achieving further gas exchanges during drought conditions, but demonstrates on the contrary, that plants have to limit water potential drops as confirmed by the modelling analysis (Figure 2b).

The relative consistency of *Ψ*_close_ among plants may appear contradictory with the large variations in minimum water potential reported by different studies[6,7,38]. However, this may highlight the importance of accounting for the multiple traits driving the demand for water when stomata are close, if we want to represent water potential decline and thus, plant dehydration. For instance, the minimum conductance (i.e. when stomata are closed) or the leaf area must be important traits driving plant water potential decline. The hydraulic model presented is consistent with this view. Accordingly, model simulations indicated that there are two main stages defining the temporal sequence leading to plant dehydration in situations of water scarcity (Figure 3). The first step is defined by the time between the start of water shortage and stomatal closure. Its duration depends principally on the rate of water uptake, given the relative constancy of *Ψ*_close_ in plants and the competition between plants for water in community ecosystems. The second stage is defined by the time between stomatal closure and plant death (100% embolism). The duration of this stage depends on a set of drought resistance traits allowing plant tissues to retain water under very high tension, to decrease water loss when the stomata are closed and to limit the decrease in water potential during embolism through deeper rooting or the release of water for internal stores. It remains to be seen how these other different traits covary with embolism resistance, are coordinated and have coevolved in plants to shape the spectrum of drought adaptation strategies.

Overall, the model analysis presented in this study, demonstrates that multiple measurable drought resistance traits can be integrated into a consistent and thermodynamically reliable formal framework to define hydraulic failure. This modelling approach must be validated carefully against temporal dynamics of water potential, hydraulic conductance, proper embolism data and experimental and field mortality for different species. However, it constitutes an important step towards assessing the consequences of drought in land plants and the effects of climate change on terrestrial ecosystem functions. It could also be a powerful tool for taking multiple traits into account in breeding strategies.

## MATERIALS AND METHODS

### 1 Data Meta-analysis

In this study we compiled embolism resistance traits and stomatal regulation traits from different species. We first compiled *Ψ*_12_, *Ψ*_50_ and slope values derived from stem vulnerability curves (i.e. the curve that relates the percent loss of conductivity to the xylem water potential) published by us during the past 20 years, representing 150 species from different biomes. Recent direct observations of embolism formation by mean of X-ray tomography confirmed the reliability of these values [21–23,40]. All stem vulnerability curves were fitted with a sigmoidal function [41]:

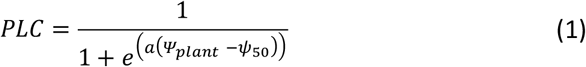

Where *PLC* is the percent loss of embolism *Ψ* is the xylem water potential, *Ψ*_50_ is the water potential causing 50% loss of plant hydraulic conductivity and *a* is a shape parameter related to the rate of embolism spread per water potential drop. Equation 1 allows computing the 2 other parameters used in this study (*Ψ*_12_, *Slope*). A more intuitive way to represent *a* is to relate it to the derivative of the function at the inflexion point of the VC, in other word, the *slope* of the linear portion of the VC:

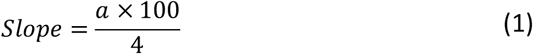

Where *slope* is expressed in %.MPa^−1^. From the *slope* and the *Ψ*_50_, the water potential at the onset of embolism, as defined by the xylem water pressure causing 12% loss of embolism, can also be computed:

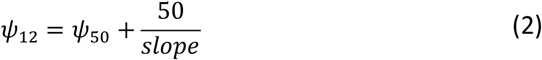

We managed to reassemble *Ψ*_50_, *Ψ*_12_ and *slope* parameters for 150 species (see Table S1). We did not considered root and leaf vulnerability curves, because it is still unclear what mechanism is responsible for the decline in hydraulic conductance measured with classical methods on these organs [42–45]. Therefore we focused our study on stem embolism that we consider being the main mechanisms responsible for extreme-drought induced mortality. We therefore neglected possible variations in embolism resistance among plant organs as we still don’t know how general is this mechanism [20,44,46].

For all the species with available stem embolism resistance traits, we collected different traits indicating the level of plant water deficit (*Ψ*) causing most stomatal closure (called *Ψ*_close_ in the main paper and hereafter). A first group of indicators was derived directly from gas exchange or transpiration measurements along with water potential data. A second indicator of *Ψ*_close_ was the bulk leaf water potential causing turgor loss [27,47,48] that was derived from pressure volume curves or from osmotic pressure at full turgor.

We searched the litterature for concurrent measurements of stomata conductance (*g*_s_) and leaf (or xylem) water potential to build g_s_(*Ψ*) curves, following the approach of [8]. We then computed the water potential corresponding to 90% of stomatal closure (*Ψ*_90gs_). Most of our *g*_s_(*Ψ*) were based on diurnal dynamic of leaf water potential of *g*_s_ and *Ψ*_leaf_ or *Ψ*_xylem_ or measurements over a drought period. In a few cases, however, we used data of concurrent measurements of water potential (leaf or xylem) and transpiration assessed through gravimetric methods (*i.e.* mass loss on detached leaves [49] or on potted plants [50] over a dehydration period obtained under constant relative humidity. To select the relevent literature, we primarily used the references provided in [8]. For each species individually, we also perform a google scholar search by using the key words “stomatal conductance” or “transpiration” AND “water potential” or “drought” or “water deficit” and the Latin name of the given species. We obtained in total, 66 species for this trait. We then searched the litterature for *Ψ*_tlp_ values. Most of them were derived from pressure volume curves [51,52], but for 10 species for which we no pressure volume curves available were found, we computed *Ψ*_tlp_ from the osmotic potential at full turgor (***π***_0_) using a linear relationship between ***π***_0_ and *Ψ_tlp_* following [53,54]. Overall, 40% of *Ψ*_tlp_ data (48 over 101 species) came from a previously published database[52], and the rest was collected from different published literature. We searched these data in google scholar, for each species for which no data were available from[52], we used the key words “osmotic potential” or “pressure volume curves” or “turgor loss point” AND the Latin name of the species. When only ***π***_0_ was measured at different time of the season only the driest date (i.e the lowest value of π_0_) was retained.

We studied the statistical associations between the different traits by using R (version 3.3.1), following a two-step procedure. First, we fitted a segmented regression to the scatter plot of *Ψ*_close_ (or its component *Ψ*_tlp_ or *Ψ*_gs90_) versus embolism resistance (*Ψ*_50_ or *Ψ*_12_) by using the package *segmented*. Then we identified (i) the break points in the *x* axis (i.e. the embolism resistance value at which there is a change in the co-variation between *Ψ*_close_ and embolism resistance) and the *y* axis intercept for this break point (i.e. the global limit for *Ψ*_close_). Second, we computed the correlation value as well as the linear regression between *Ψ*_close_ and embolism resistance for the data on either side of the break point. In addition to the results developed in the main manuscript, we provided a separate analysis per group (gymnosperm and angiosperm) and per trait (*Ψ*_50_, *Ψ*_12_, *Ψ*_gs90_, *Ψ*_tlp_) in Table S2. All parameters used in this study are given in a supplementary Excel file.

### 2 Model: description, simulation and validation

We used a simplify discrete-time-hydraulic-model (called Sur_Eau) to simulate the time until hydraulic failure for the spectrum of embolism resistance reported in our database, and according to different hypothesis regarding the stomatal regulation of transpiration. Sur_Eau relies on the principle of the original Sperry's model [19] but has been simplified to consider only two resistances (rhizosphere and plant). This simplicity makes easier its applicability with only one stem VC and avoids making assumptions on hydraulic segmentation, a phenomenon that depends on mechanisms that are still controversial.

#### 2.1 *Description of the Sur_Eau model*

Sur_Eau assumes that liquid water flow through the soil-plant system is exactly compensated by gaseous water losses at the plant foliage surface (i.e. steady state condition) which is true at large time steps (>1day) or for small plants. The assumption is also made that leaf and air temperatures are roughly equal, which is reasonable in well coupled canopies. We can then write:

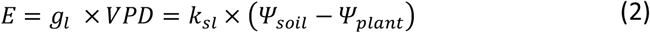

where *g*_l_ is the leaf conductance to vapor water, VPD is the vapor pressure deficit at the leaf surface, *Ψ*_soil_ is the soil water potential, *Ψ*_plant_ is plant water potential, and *K_sl_* is the plant leaf area specific hydraulic conductance over the soil to leaf pathway. *g*_l_ includes both the stomatal, cuticular and boundary layer conductances of the leaf. The control of E through stomata has been treated through different assumptions that are described below (Appendix 3). *k*_sl_ was computed as the result of two conductances in series:

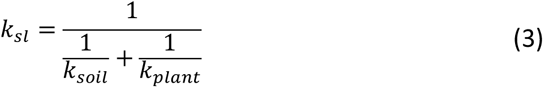

where *k_soil_* is the hydraulic conductance of the soil to root surface pathway and *k_plant_* the hydraulic conductance of the whole plant (i.e. from the roots to the leaves).

*K_plant_* was allowed to vary only to account for loss of hydraulic conductivity caused by xylem embolism [55]:

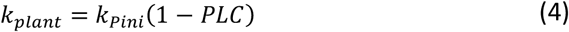

where *k_Pini_* is the initial (*i.e* pre-drought) plant hydraulic conductance, and PLC is the percent loss of plant hydraulic conductance due to xylem embolism. PLC is computed at each time step by using the sigmoidal function for the vulnerability curve (VC) to embolism (see equations 1 to 3).

The model considers the capacitive effect of xylem embolism and symplasm dehydration. The water freed by air filling of the *apoplastic* reservoir feeds the water stream of the system and thus dampen the water potential decrease [56]. Following^24^, we considered that any change in PLC is followed by a proportional change of the water volume that is freed back to the system:

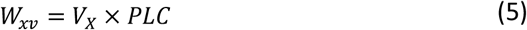

Where *W*_xv_ is the amount of water freed to the system and *V_x_* is the total water filled xylem volume of the plant (m^3^) and PLC is defined in Equation 1. *V_x_* was computed as:

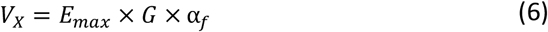

Where *E_max_* is the maximum diurnal transpiration, *α_f_* is the apoplasmic fraction of the plant and *G* is the ratio of the total amount of water in the xylem conduits of a tree (*i.e.* apoplasmic volume) to the maximum diurnal transpiration. The capacitive effect of cavitation on plant survival is mainly sensitive to *G* factor and its range of variation. The effect of G on cavitation dynamics is discussed in^24^. We also accounted for the capacitive effect of symplasm dehydration (*i.e.* the water released by the symplasmic tissue *W_sv_*) by using the same formulation as for cavitation (Equation 6). Computations were based on the symplasmic fraction (1− *α_f_*) of the plant and PLC was replaced by the relative water content of the symplasm (*R*_s_). *R*_s_ was computed from *Ψ*_leaf_ by using the pressure volume curve equations (Appendix 3).

The variations of the soil and the rhizophere conductance (*k*_soil_), as well as the mean soil water potential in the root zone are computed with Van-Genuchten-Mualem equations [33,57] from the unsaturated hydraulic conductivity of the soil, scaled to the rhizophere according to the Gardner-Cowan formulation [58,59]. The rhizophere conductance (*k*_soil_) can be expressed as:

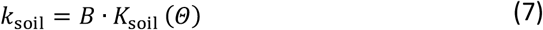

where *K*_soil_ is the unsaturated hydraulic conductivity of the soil at given water content (*Θ*) or water potential (see below), and *B* is the root density conductance factor that accounts for the length and geometry of the root system. *B* is based on the implicit assumption of a uniform roots distribution in a soil layer following the Gardner-Cowan formulation^28^. *B* is also called the “single root” approach [60] as it is equivalent to assuming that plant water uptake occurs from a unique cylindrical root that has access to a surrounding cylinder of soil:

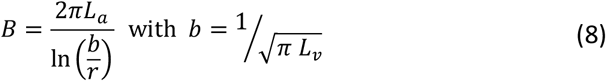

where *L_a_* is the root length per unit area, *r* is the mean root radius, and *b* is the half of mean distance between neighboring roots. *b* can be evaluated from *L_v_*, the root length per unit soil volume. *k*_soil_ decreases with decreasing *Ψ*_soil_ because of the displacement of water filled pores by air, as capillary forces linking water to soils particles fail with increasing tension, thus creating dry non-conductive zones in the rhizosphere. The parametric formulation of^25^ for the water retention curve was used in combination with the equation of Mualem (1976)^26^ to compute *Ψ*_soil_ and the unsaturated hydraulic conductivity of the soil (*K*_soil_) as a function of soil relative extractable water content (*Θ*)^25^. The formulation of *Ψ*_soil_(Θ) is as follows:

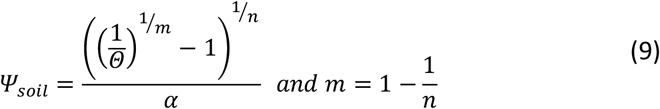

where *m*, *n* and *α* are empirical parameters describing the typical sigmoidal shape of the function. Mualem (1976) provides the formulation for the evolution if hydraulic conductivity with soil water content *k*_soil_(Θ):

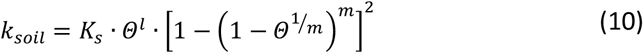

where *K*_s_ is the saturated hydraulic conductivity, *l* is a parameter describing the pore structure of the material (usually set to 0.5), and *m* is again fixed as in Equation 11. The relative extractable water content (Θ) is expressed as a function of volumetric soil water content as follows:

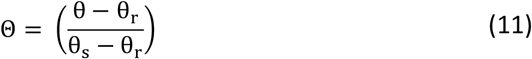

where θ, θ_s_ and θ_r_ are the actual relative soil water content and the relative soil water content at saturation and at wilting point respectively. θ_s_ and Θ_r_ are parameters measured in laboratory or derived from soil surveys using pedotransfert functions. By contrast, θ is a variable, dynamically changing with changes of the soil water reserve (WR). The water reserve depends on the soil volume θ_s_ and Θ_r_. The calibration and sensitivity analysis for all parameters is provided below (Appendix4).

#### 2.2 *Dynamic simulations*

Dynamic simulations of the different variables were performed by the mean of the discrete time model described above. Under well-watered conditions, transpiration (E) is forced at a constant value, assuming a constant high vapor pressure deficit. Then, *E* is regulated with decreasing water potential as leaves loose turgor inducing stomata closure, with a submodule based on the pressure volume curve (following [32]; see Appendix 3). At each time step the soil water reserve (WR) is first computed and then used to compute all other variables. WR is then computed as a result of a water balance:

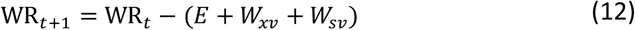

Where *E* is the cumulated transpiration over the time step, *W_xv_* is water release due to cavitation and *W_sv_* is the water release due to symplasm dehydration (Equation 7 & 8 and the following text). The time step was set to 0.1 day, but increasing this value up to 0.5 or down to lower values had little influence on the general pattern of our results. *E* was computed as follows:

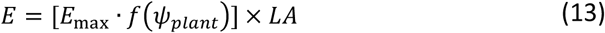

where *E*_max_ is the maximal rate of transpiration, LA is the plant leaf area and f(*Ψ*_plant_) represents the stomatal regulation which was set according to different hypothesis as described in the next section (Appendix 2). The calibration for *E*_max_ and LA as well as all other parameters are provided in Appendix 4.

#### 2.3 *Hypothesis testing on the interrelation between Ψ_close_ and Ψ_50_*

We used the above described model to evaluate the role of *Ψ*_close_ and *Ψ*_50_ in determining the survival time until hydraulic failure under drought for the full range of embolism resistance reported in the database. Three different hypotheses regarding how stomata regulate transpiration were tested (Figure 2A): (1) No stomatal regulation of *E* (*Ψ*_close_=-∞) whatever the embolism resistance, (2) *E* regulation to maintain *Ψ*_plant_ above the water potential causing the onset of embolism (*Ψ*_plant_ >*Ψ*_12_) and (3) Early stomatal regulation of *E* during drought, so that *Ψ*_close_ varied with *Ψ*_12_ until *Ψ*_close_=-3 MPa, in accordance with the global limit observed in empirical data (Figure 1c, main manuscript). For each of these hypothesis, survival time was computed for the full range of embolism resistance encountered in our database, from *Ψ*_50_= −1.5 to *Ψ*_50_=-15 MPa, every 1MPa step. A more detailed analysis of these simulations is given in the Appendix 2.

#### 2.4 *Model validation: survival time during drought from drought mortality experiments*

To validate the relationship between embolism resistance traits (*Ψ*_50_) and survival time predicted by the model under different hypothesis, with used data from drought mortality experiments. We found survival time data for 15 species covering a wide range of embolism resistance (*Ψ*_50_ from −1.5 to −11), from four different drought mortality experiments published recently [29–31,50]. One study was performed on gymnosperm species only^17^ and three other studies were performed on angiosperm species [29–31,50]. All these experiments were made under semi-controlled conditions on potted seedlings or saplings and shoot death was recorded at different time since the beginning of an experimentally imposed drought. Here we used the average time needed to reach 50% death (T_50_) after the beginning of the drought treatment as an indicator of survival time during drought. In each experiment, identical soil volume and climate were used across species. However, there were differences in air relative humidity and soil volume across experiments (both of them can strongly affect the survival time during a water deficit episode), particularly between the gymnosperm and all angiosperm experiments. This precluded the direct comparison of the survival time across the four different studies. To overcome this problem, we standardized the T_50_ of each experiment by differences in soil volume or air relative humidity (Appendix1).

## Appendix and includes

**Appendix 1:** Methods for obtaining survival time during drought for model validation

**Appendix 2:** Detailed simulations of the three hypothesis tested for the stomata regulation of embolism.

**Appendix 3:** Turgor loss model and symplasm water content simulation

**Appendix 4:** Model parameterization and sensitivity analysis

## Supplementary material includes

**Table S1:** Summary description of the data collected

**Table S2**: correlation and linear regression between *Ψ*_close_, *Ψ*_tlp_, *Ψ*_gs90_ and embolism resistance (*Ψ*_12_, *Ψ*_50_) on either side of the breakpoints for all species or gymnosperm and angiosperm separately.

**Excel file of database of stomatal and stem hydraulic traits**

